## Supplementary files for "WDR82 mediated H3K4me3 is associated with tumor proliferation and therapeutic efficacy in pediatric high-grade gliomas"

**Running title:** WDR82 mediated H3K4me3 in pediatric gliomas

Nitin Wadhwani,^1^ Sonali Nayak,^2^ Yufen Wang,^3^ Rintaro Hashizume,^4^ Chunfa Jie,^5^ Barbara Mania-Farnell,^6^ Charles David James,^7^ Guifa Xi,^2,7^ Tadanori Tomita,^2,7^

1. Department of Pathology, Ann & Robert H. Lurie Children’s Hospital of Chicago, Northwestern University Feinberg School of Medicine, Chicago, IL 60611 USA
2. Division of Pediatric Neurosurgery, Ann & Robert H. Lurie Children’s Hospital of Chicago, Northwestern University Feinberg School of Medicine, Chicago, IL 60611 USA
3. Department of Radio-oncology, Northwestern University Feinberg School of Medicine, Chicago, IL 60611 USA
4. Department of Pediatrics, Northwestern University Feinberg School of Medicine, Chicago, IL 60611 USA
5. Department of Biochemistry and Nutrition, Des Moines University Medicine and Health Sciences, Des Monies, IA 50312 USA
6. Department of Biological Science, Purdue University Northwestern, Hammond, IN …USA
7. Department of Neurological Surgery, Northwestern University Feinberg School of Medicine, Chicago, IL 60611 USA

**Corresponding author**

Guifa XI, MD, PhD

The Department of Neurological Surgery, Northwestern University Feinberg School of Medicine

Address: Simpson Querrey Biomedical Center, Room 6-500

303 Superior Street, Chicago, IL 60611

Tadanori Tomita, MD

Division of Pediatric Neurosurgery, Ann & Robert H. Lurie Children’s Hospital of Chicago, Northwestern University Feinberg School of Medicine

Address: 225 E Chicago Ave, Box 28

Chicago, IL 60611

**Supplementary Materials and Methods**

**Tumor sphere formation assay.** Cells were cultured in T25 flasks with or without Dox (2µg/ml), incubated at 37°C with 5% CO_2_ for three days, harvested as single cell suspension and counted prior to plating 2000 cells/well in ultra-low attachment 24-well plates (Cat#3473, Corning Incorporated). Cell culture medium contained DMEM:F12 (1:1) (Cat#12660-012), B27 (Cat#A35828-01), EGF (Cat#PHG0311) 20ng/ml, b-FGF (Cat#PHG0021) 20ng/ml from Gibico. Cell growth was monitored daily, images were captured and the number of spheres were counted with EVOS FL microscopy (Life Technologies) at day 3.

**MTS assay for chemosensitivity assessment.** pHGG cells were used to determine if reducing WDR82 through inducible knockdown affected cell viability in the absence or presence of chemotherapeutic drugs, vincristine (VCR) and cisplatin (CDDP). For the cells in the absence of chemotherapeutic drugs, 1×10^4^ cells/100μl were plated in 96-well plates with complete cell culture medium with or without Dox (2µg/ml), and subjected to 3-(4, 5-dimethylthiazol-2-yl)-5-(3-carboxymethoxyphenyl)-2-(4-sulfophenyl)-2H-tetrazolium (MTS, Promega) assay. Cells in the presence of chemotherapeutic drugs, were harvested and counted with TC-20^TM^ Cell Counter (Bio-Rad Laboratories Inc.), equal numbers of cells were plated in 96-well plates with complete cell culture medium in the absence or presence of (Dox) (2µg/ml) for 2 days following treatment with cisplatin (CDDP, 10^-3^ to 10^3^ µM), and subjected to MTS assay after 72h.

**Clonogenic survival assay**. Cells were seeded into 6-well tissue culture plates and allowed to adhere. Attached cells were irradiated (0, 1, 2 or 4Gy) and treated with or without Dox (SKU#D9891-10G, Sigma-Aldrich) at 2µg/ml 2 hours after irradiation. Radiation was delivered by a ^137^Cs source (Mark I, model 68A irradiator, JL Shepherd & Associates) as previously described.^1^ Cells were incubated with Dox for 2 weeks, at which time colonies were stained with methylene blue (0.66% solution in 95% ethanol) and counted. Plating efficiencies were calculated as the ratio of the number of colonies formed to the number of cells seeded. Colonies of >50 cells were counted for surviving fraction determinations. Surviving fractions were calculated as the plating efficiency of treated cells divided by the plating efficiency of control cells, as previously described.^2^ Dose enhancement factors were calculated at 10% survival.

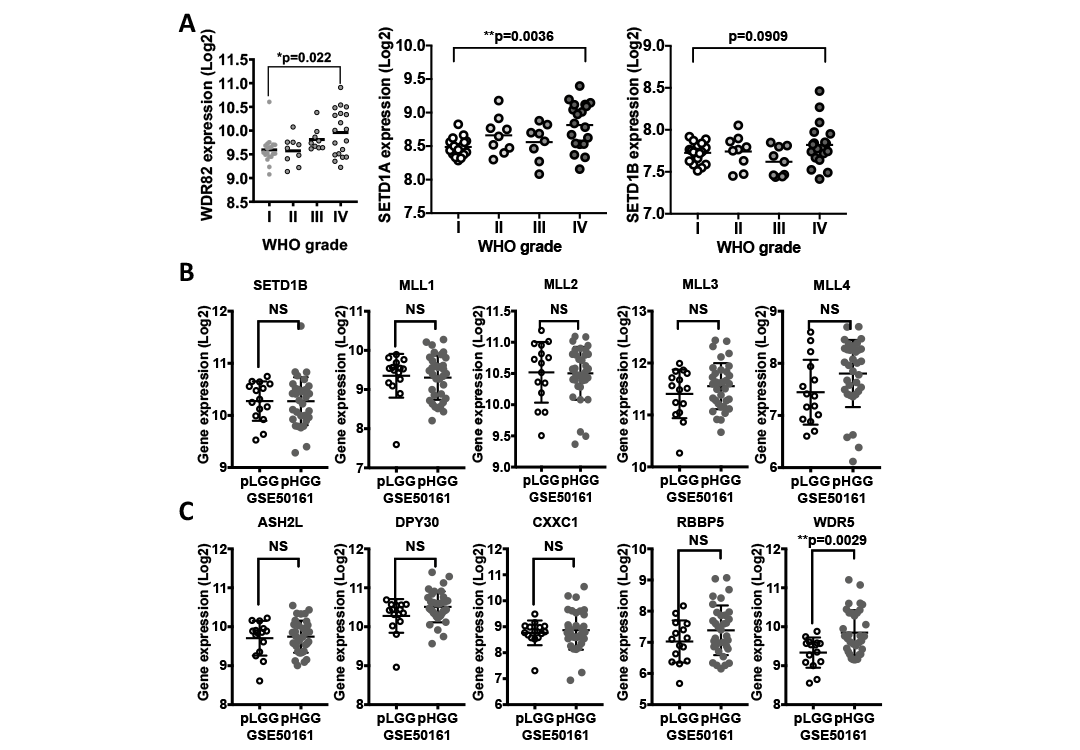

**Supplementary Figure 1. In silico analysis of expression of subunits of human SET-COMPASS complexes in pediatric gliomas. A.** Expressions of WDR82, SETD1A and SETD1B in pediatric gliomas. **B and C.** In silico analysis of GEO dataset GSE50161 showing expression of SETD1B and MLL1-4 (B) and expression of SETD1A/B subunits in pediatric low- (pLGG) and high- (pHGG) grade gliomas (C).

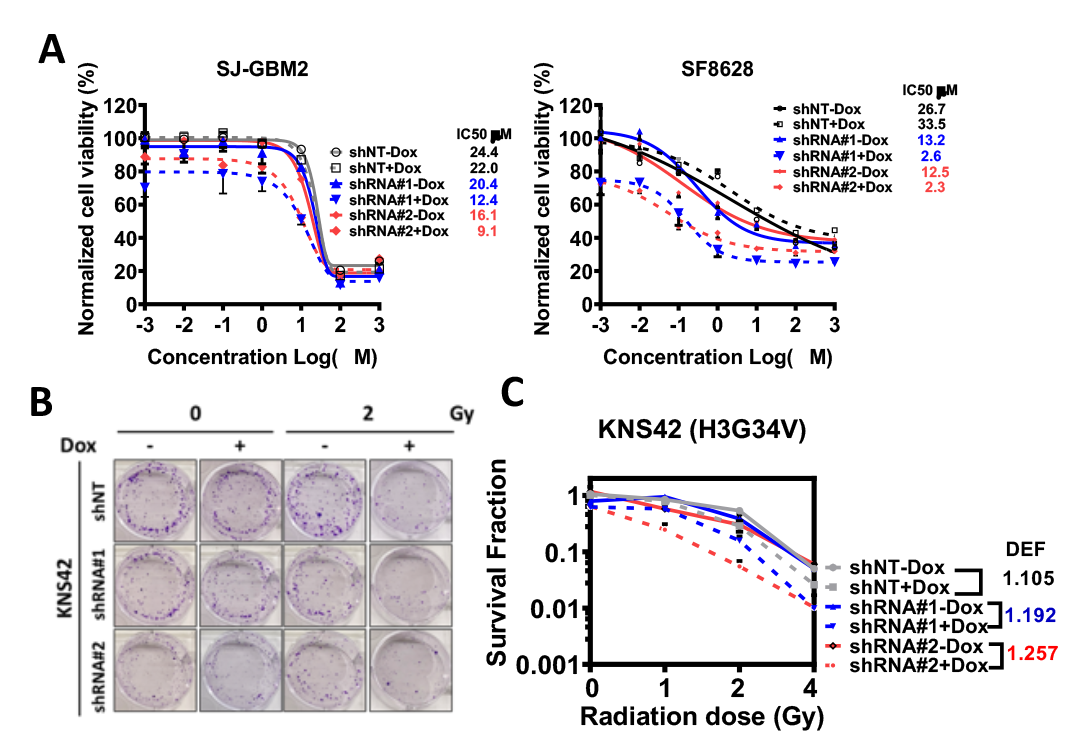

**Supplementary Figure 2. WDR82 knockdown increases chemotherapeutic and radiation sensitivity.** A. Normalized cell viability of SJ-GBM2 and SF8628 cells transduced with inducible shRNAs #1 and #2 against WDR82 and control non-target shRNA (shNT) in the absence or presence of Dox, 2μg/ml, following treatment with cisplatin (CDDP). B. Representative images show colonies from 1000 cells in the absence (-) or presence (+) of Dox, following radiation therapy. (C) Quantitative results from B.

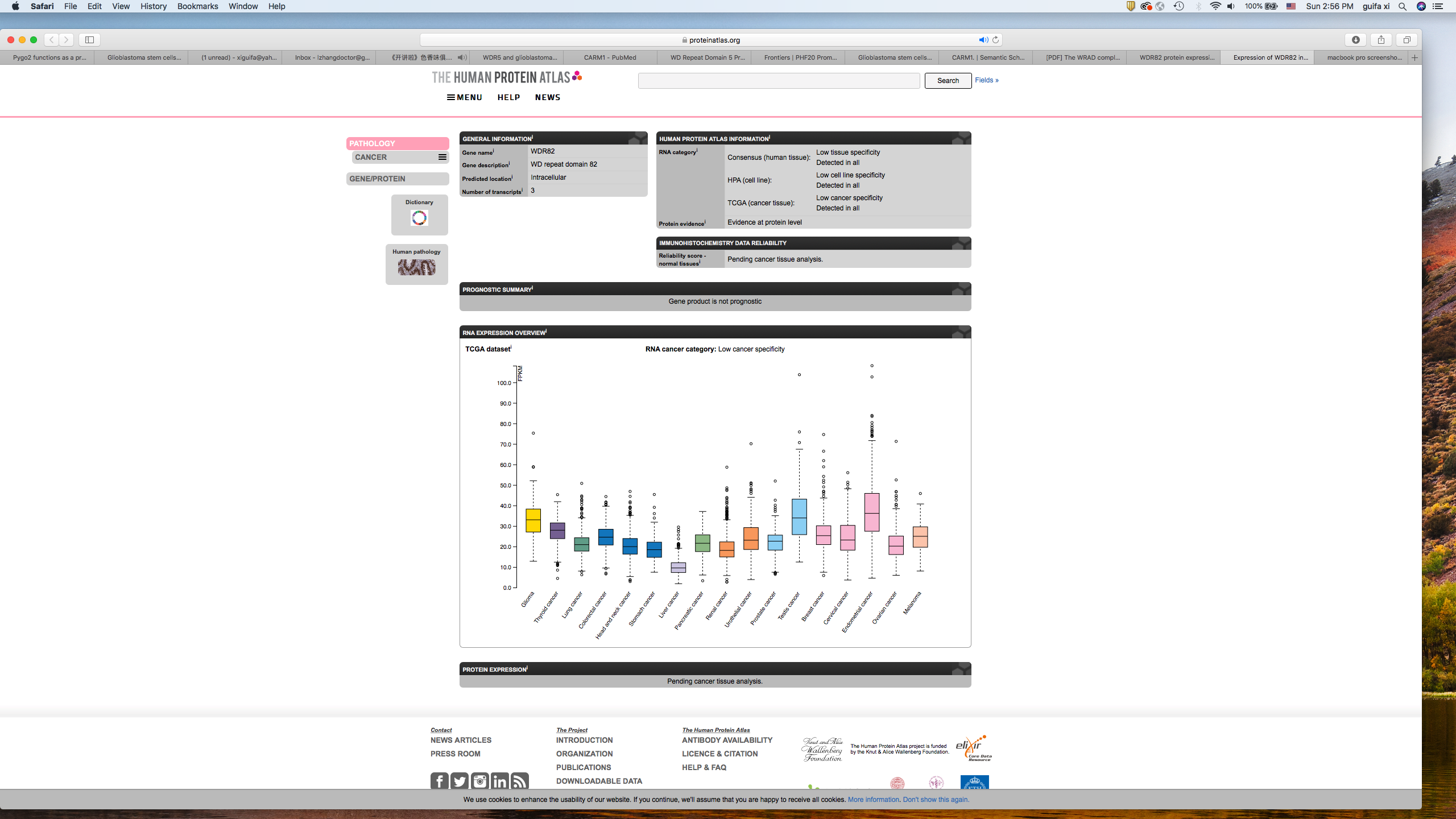

**Supplementary Figure 3. WDR82 expression in human cancers presented with TCGA dataset** (https://www.proteinatlas.org/ENSG00000164091-WDR82/pathology).

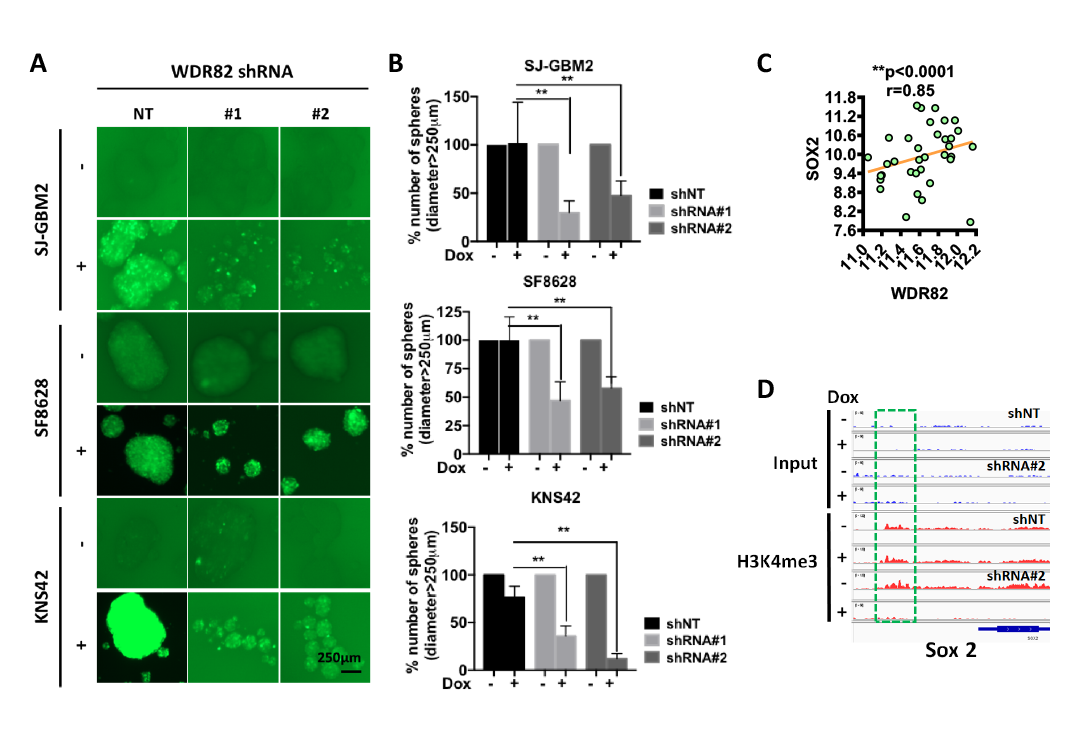

**Supplementary Figure 4. WDR82 is associated with stem cell characteristics.** Representative images (**A**) and normalized quantitative results for sphere diameter > 250µm (**B**), show tumor sphere formation in SJ-GBM2 (H3WT), SF8628 (H3K27M) and KNS42 (H3G34V) pHGGs transduced with control shNT, or shRNA#1, or shRNA#2 against WDR82, in the absence (-) and presence (+) of Dox (2µg/ml). **C**. Correlation between WDR82 and SOX2 through in silico analysis of dataset GSE50161. (**D**) ChIP seq results from SJ-GBM2 cells show H3K4me3 level at the SOX2 promoter. SJ-GBM cells non-treated or treated withDox, 2µg/ml.

**Supplementary Table 1** Clinicopathological information for pediatric primary gliomas IHC stained for H3K4me3.

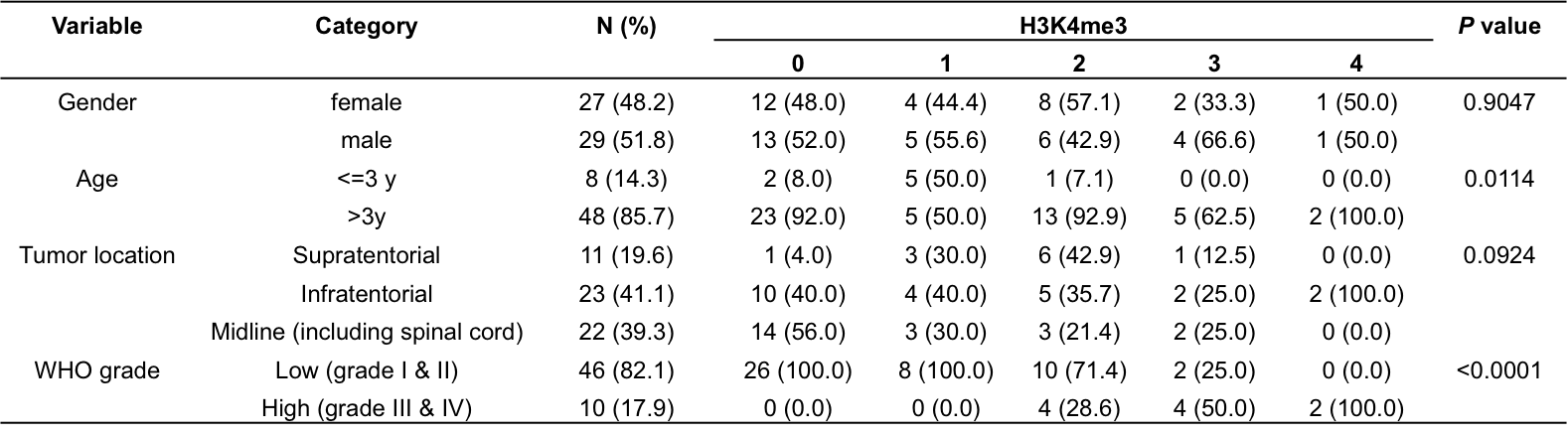

*P* value was calculated by Fisher's Exact test.
